## Supplementary Figures for "Biologically-informed self-supervised learning for segmentation of subcellular spatial transcriptomics data"

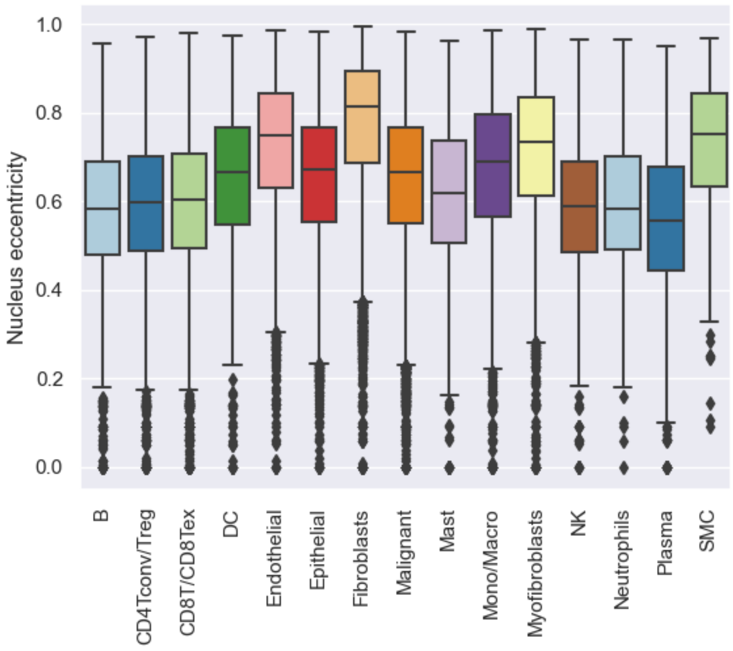


Supplementary Figure 1: Boxplots showing the eccentricity of nuclei for different cell types for Xenium-BreastCancer1.

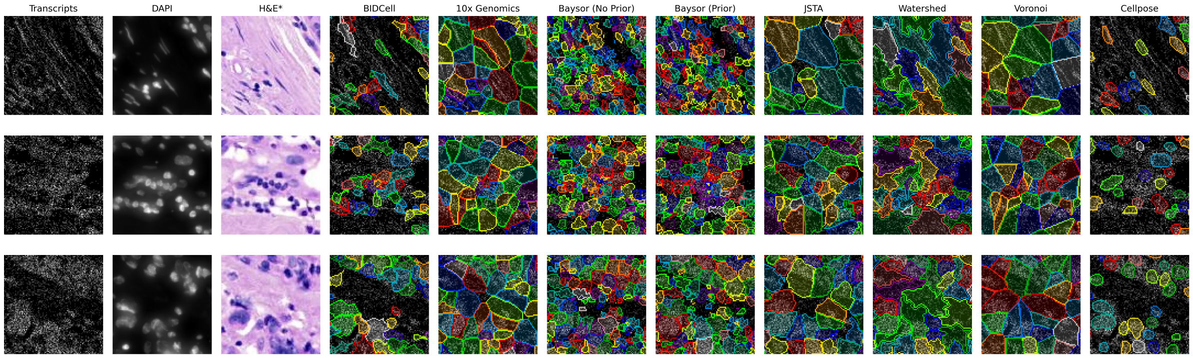


(a) Xenium-BreastCancer1

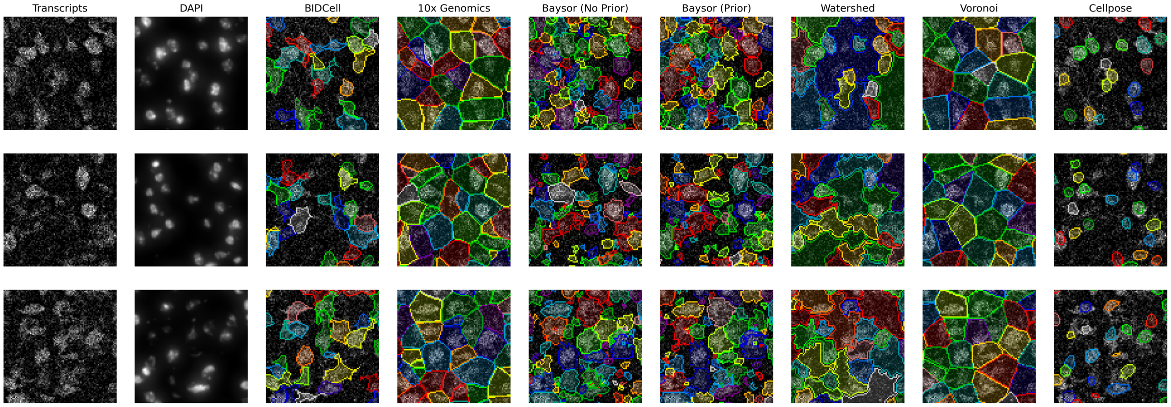


(b) Xenium-MouseBrain

Supplementary Figure 2: Further selective illustration of BIDCell and other segmentation methods. H&E images are shown for visualisation purposes only. BIDCell generate morphologies that exhibit better visual correspondence with the input images.

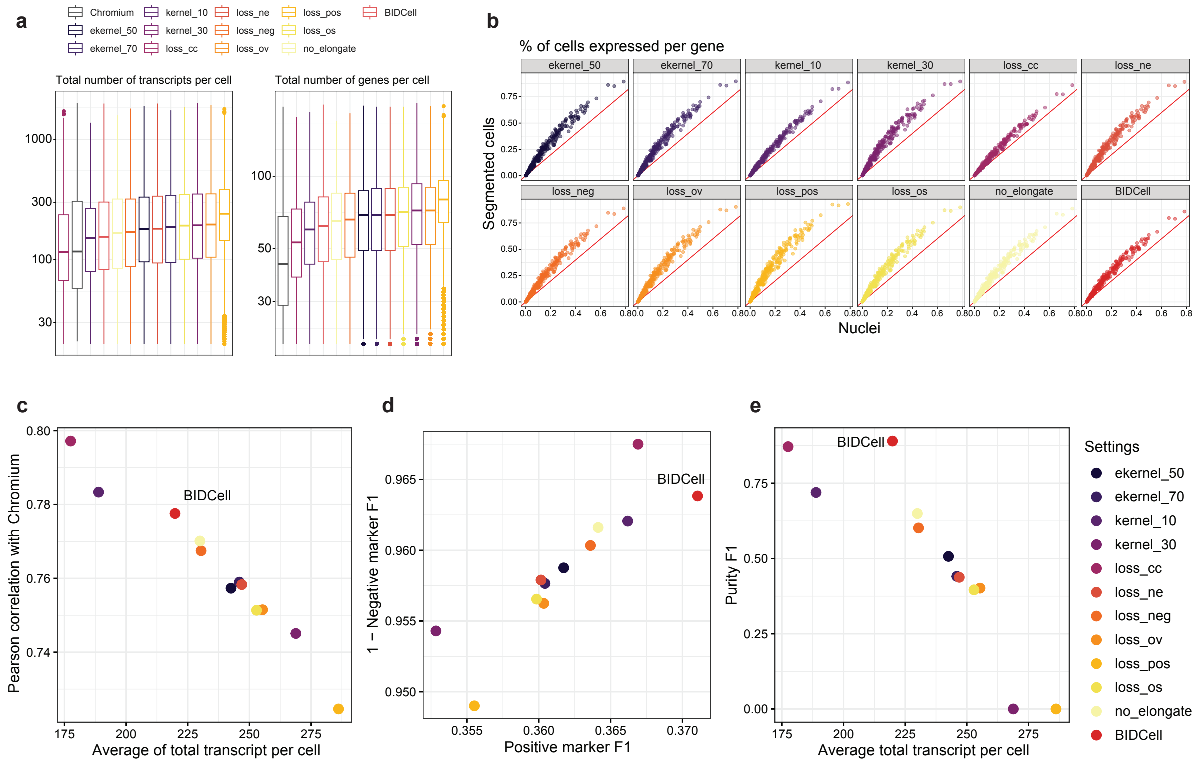


Supplementary Figure 3: Ablation study of different settings of main parameters: `ekernel_50` and `ekernel_70`, where $l_t$ of the elliptical kernel in the cell-calling loss was 50 or 70; `kernel_10` and `kernel_30`, where the diameter of the circular kernel in the cell-calling loss was 10 and 30; `loss_`, where each loss was individually set to zero; `no_elongate`, where all cells were assumed a circular shape; and `BIDCell`, the proposed settings. `BIDCell` achieved the highest purity, with more transcripts captured and a relatively high Pearson correlation compared to `loss_cc`.

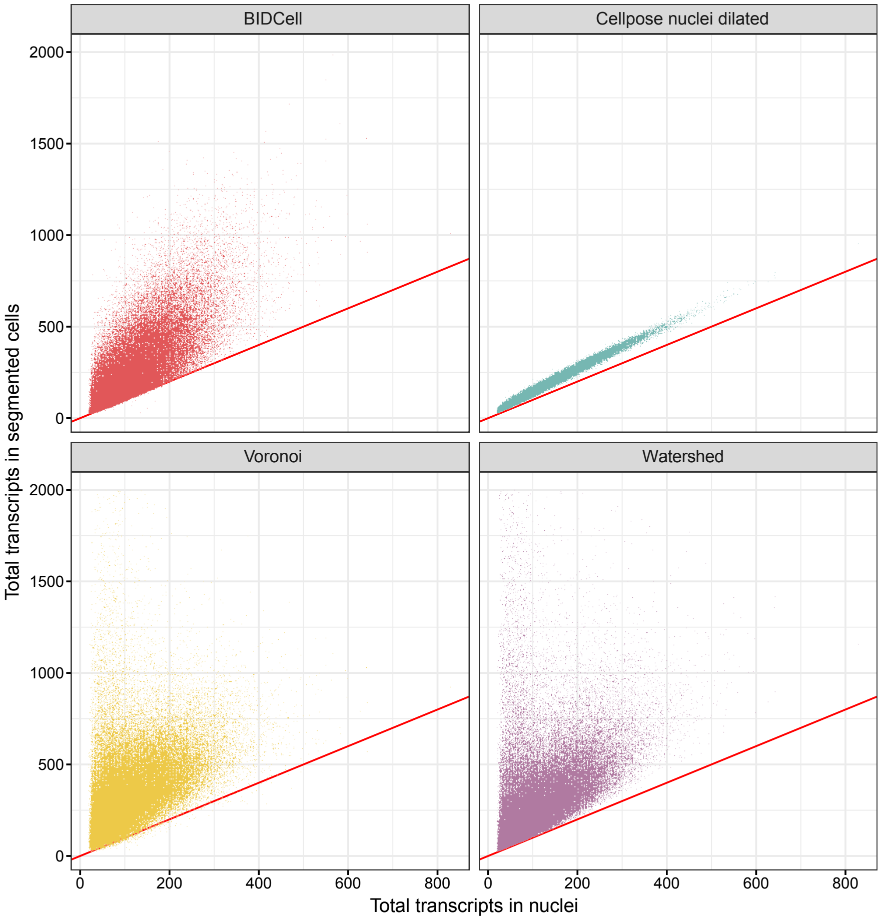


Supplementary Figure 4: Total transcripts per cell for Xenium-BreastCancer1, comparing between Cellpose nuclei and cells segmented using four methods (BIDCell, Cellpose nuclei dilated, Voronoi, and Watershed). The red diagonal line indicates where the total number of transcripts in the nuclei is equal to the number of transcripts in the segmented cells.

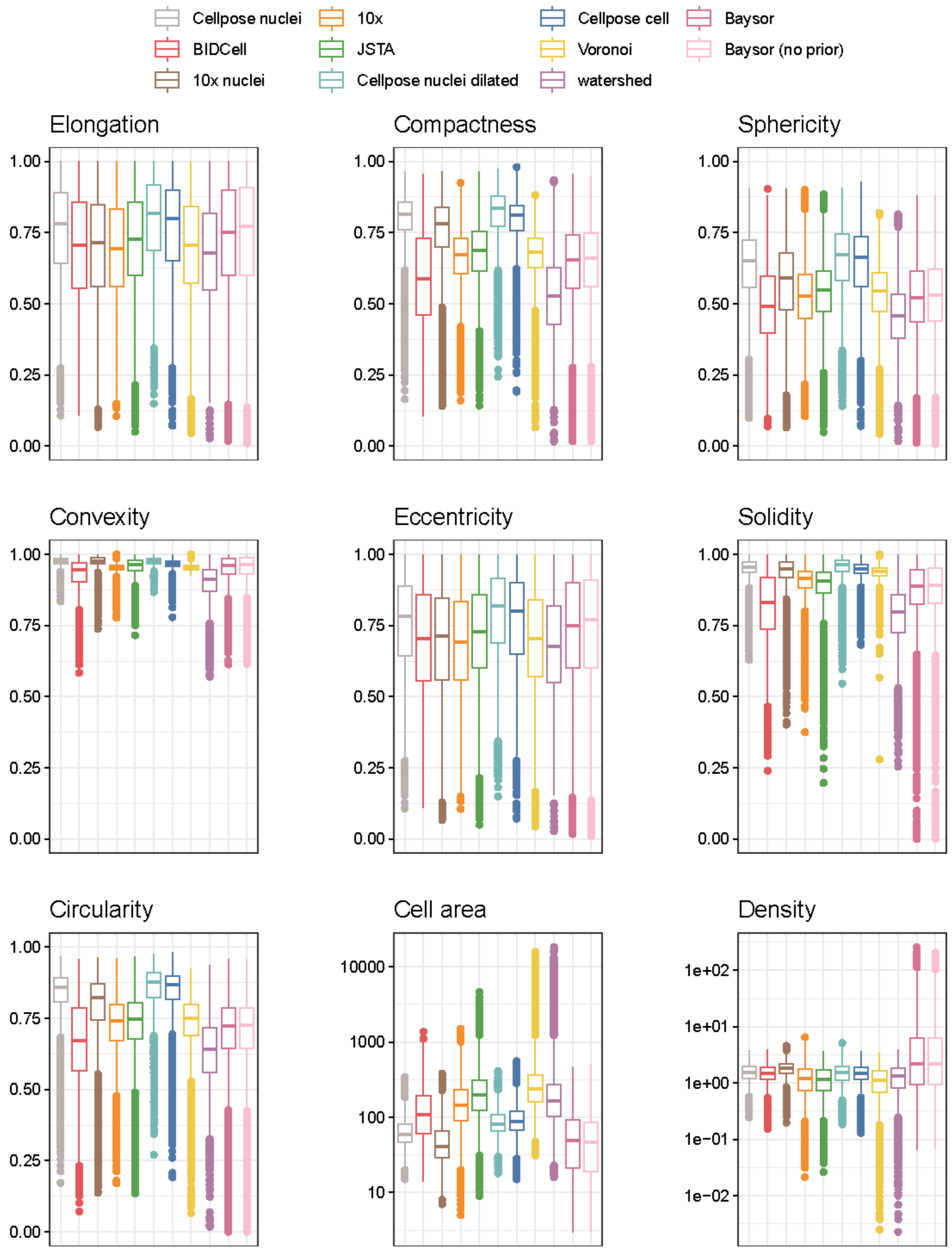


Supplementary Figure 5: Cell morphology metrics (elongation, compactness, sphericity, convexity, eccentricity, solidity, circularity, cell area, and density) of Xenium-BreastCancer1 for different cell segmentation methods. Each box indicates results for one method.

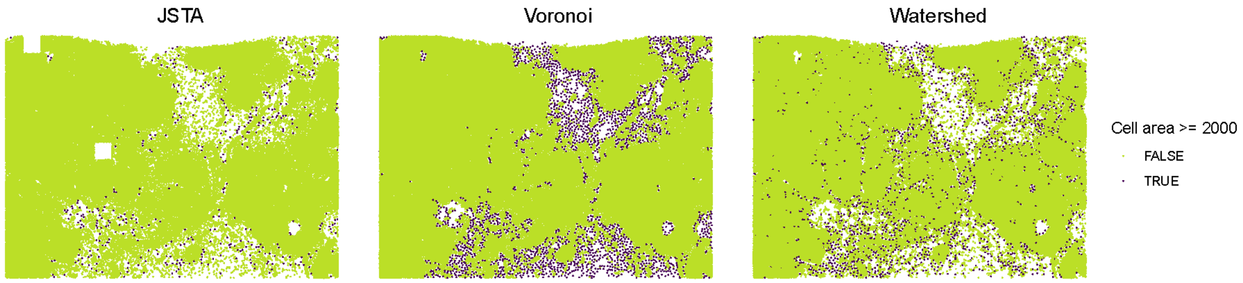


Supplementary Figure 6: Cell area outlier in three methods (JSTA, Voronoi, and Watershed), where we observe that these methods tend to produce cells with large areas in the sparse regions.

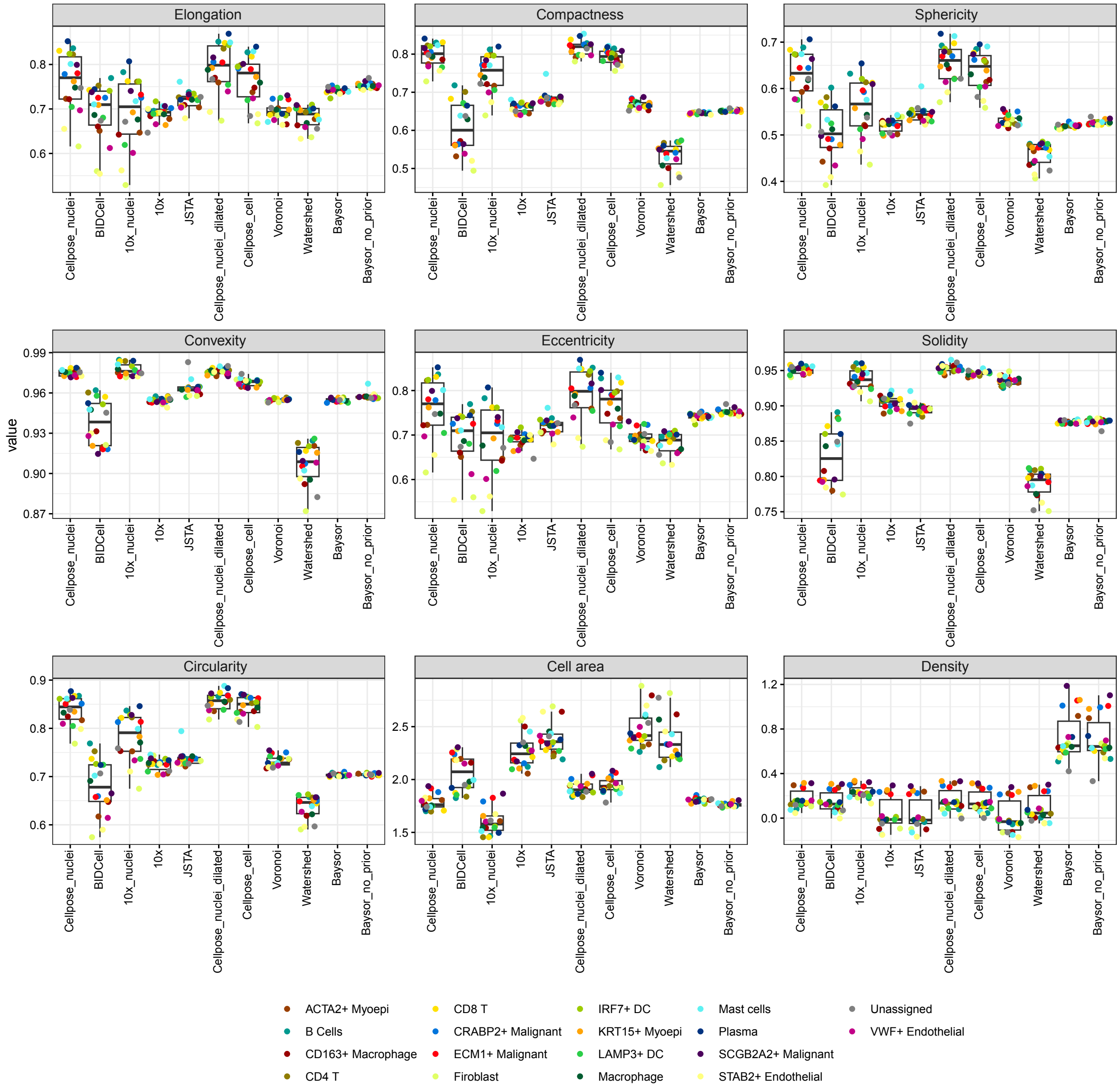


Supplementary Figure 7: Boxplots of average cell morphology metrics (elongation, compactness, sphericity, convexity, eccentricity, solidity, circularity, cell area, and density) per cell type of Xenium-BreastCancer1 for different cell segmentation methods, where each point indicates one cell type.

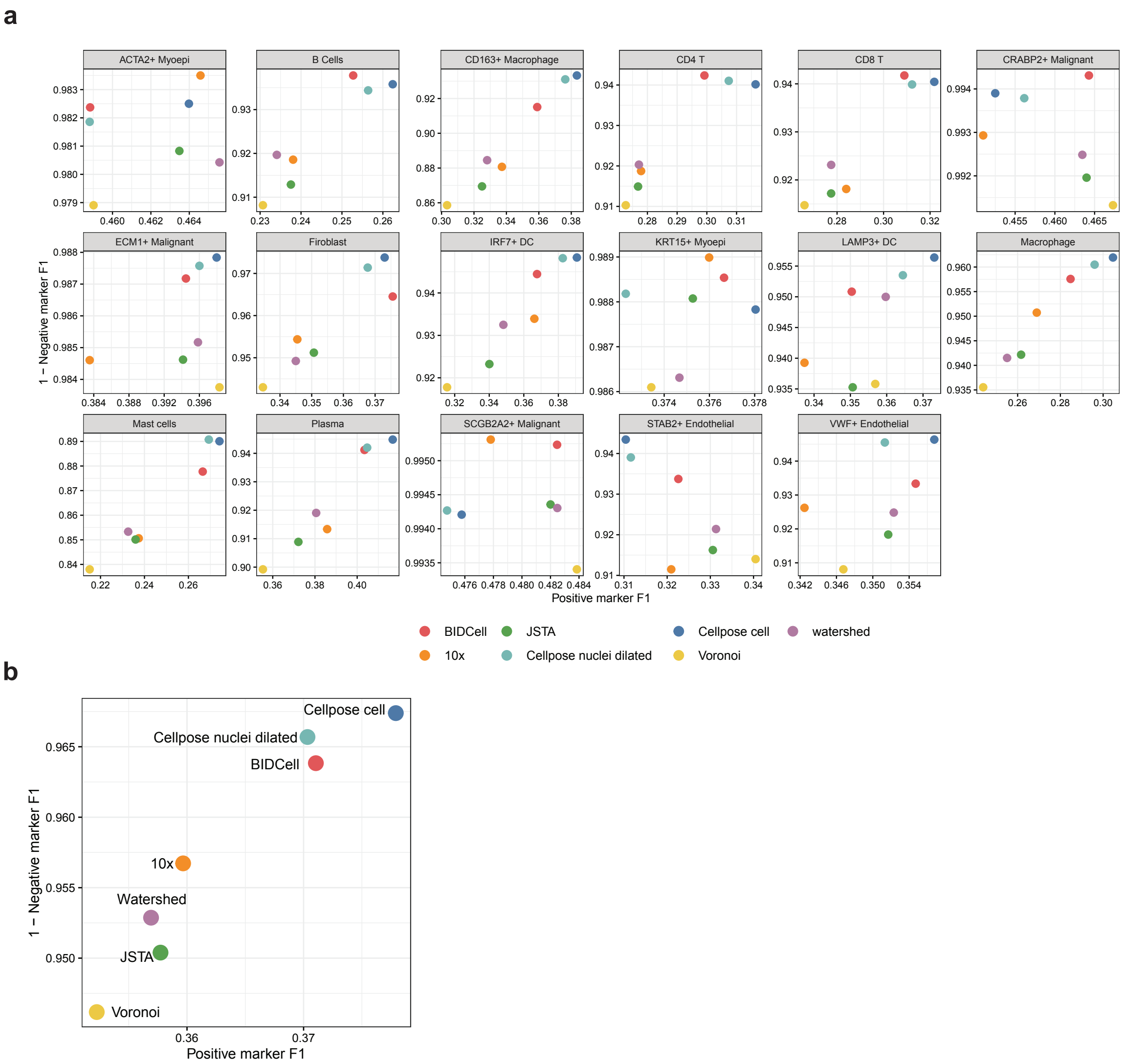


Supplementary Figure 8: Expression purity benchmarking results for Xenium-BreastCancer1. (a) Scatter plots showing the positive marker F1 scores vs. the 1 - negative marker F1 scores for each of the cell type, where each dot is one method. (b) Scatter plots showing the overall positive marker F1 vs. the 1 - negative marker F1.

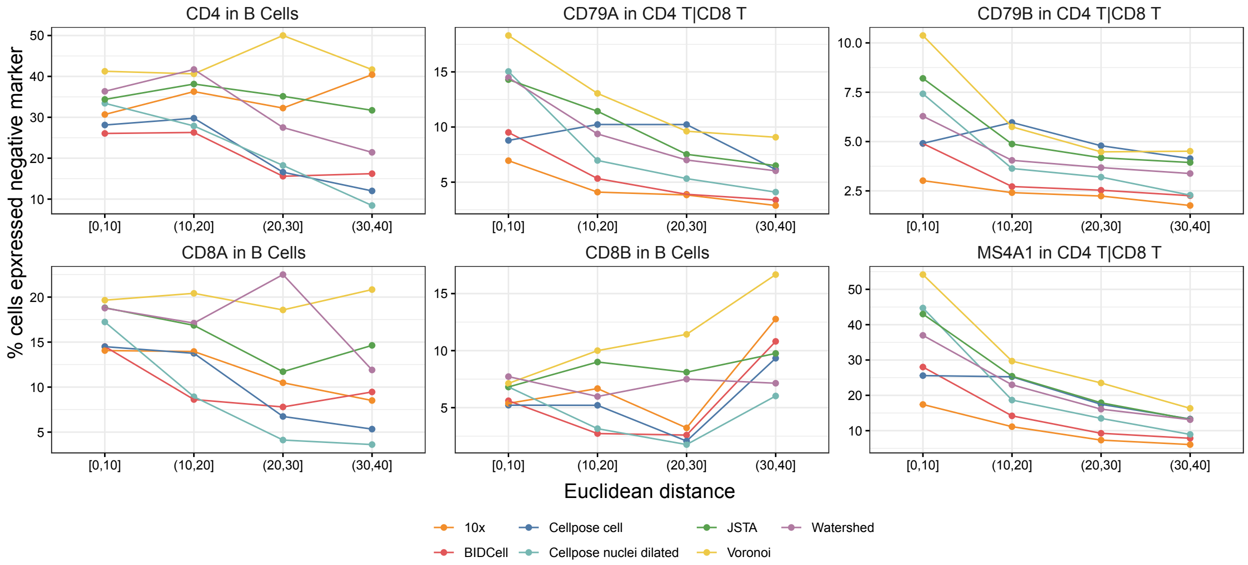


Supplementary Figure 9: Neighbouring contamination results for Xenium-BreastCancer1. The line plots indicate the percentage of B cells expressing the unwanted T cell marker CD4, CD8A, and CD8B against its distance from the nearest T cell, where the cells are grouped by a certain distance range; and the percentage of T cells expressing CD79A, CD79B, and MS4A1, against the distance from the nearest B cells. A lower percentage is better, and each line represents a different method.

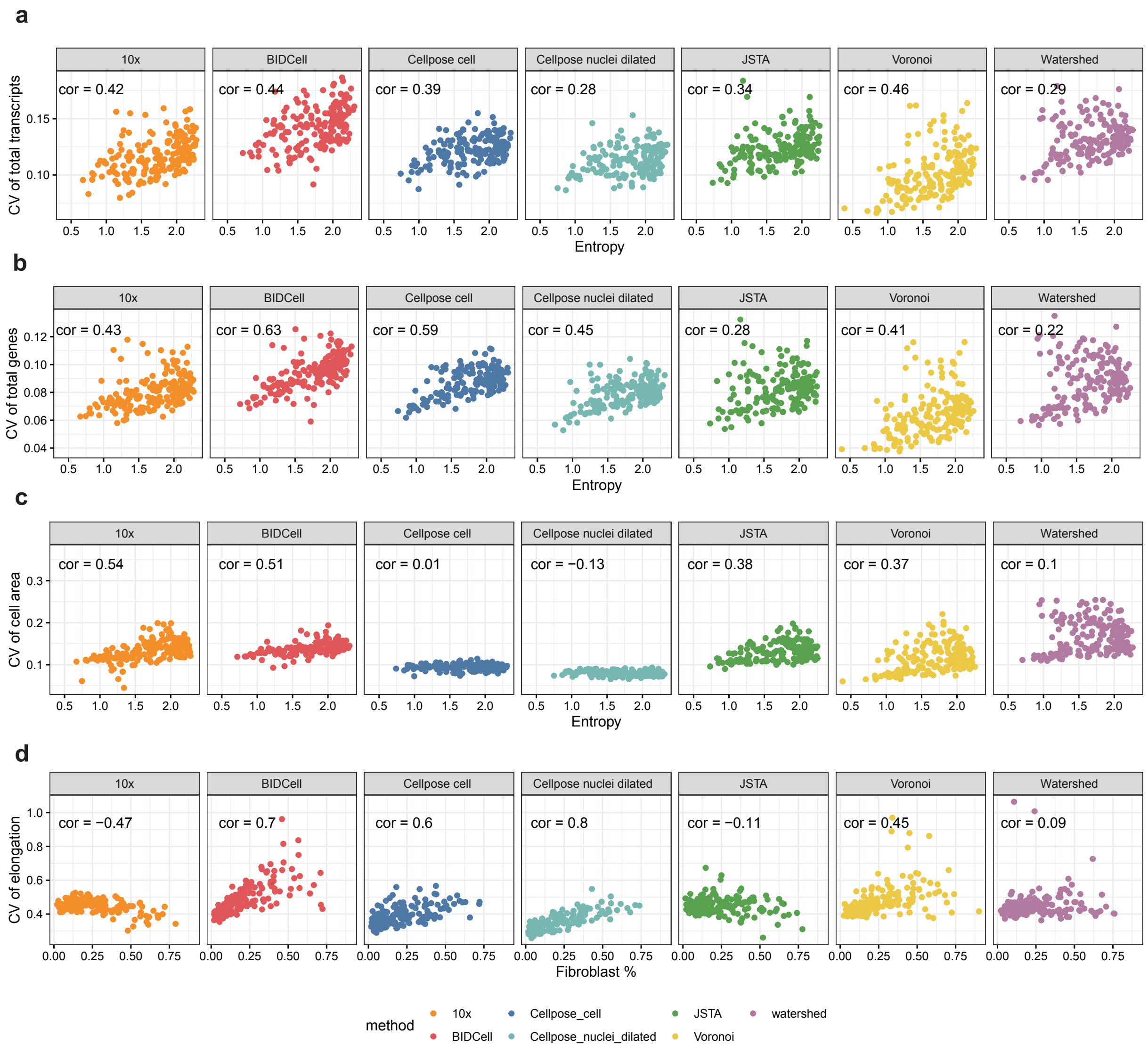


Supplementary Figure 10: Spatial diversity results for Xenium-BreastCancer1, corresponding to Figure 3g-h. Scatter plots showing the association between cell type entropy and (a) coefficient of variation of the total transcripts; (b) coefficient of variation of the total genes; and (c) coefficient of variation of cell area. (d) Scatter plots showing the association between the coefficient of variation of elongation and proportion of fibroblasts in the data.

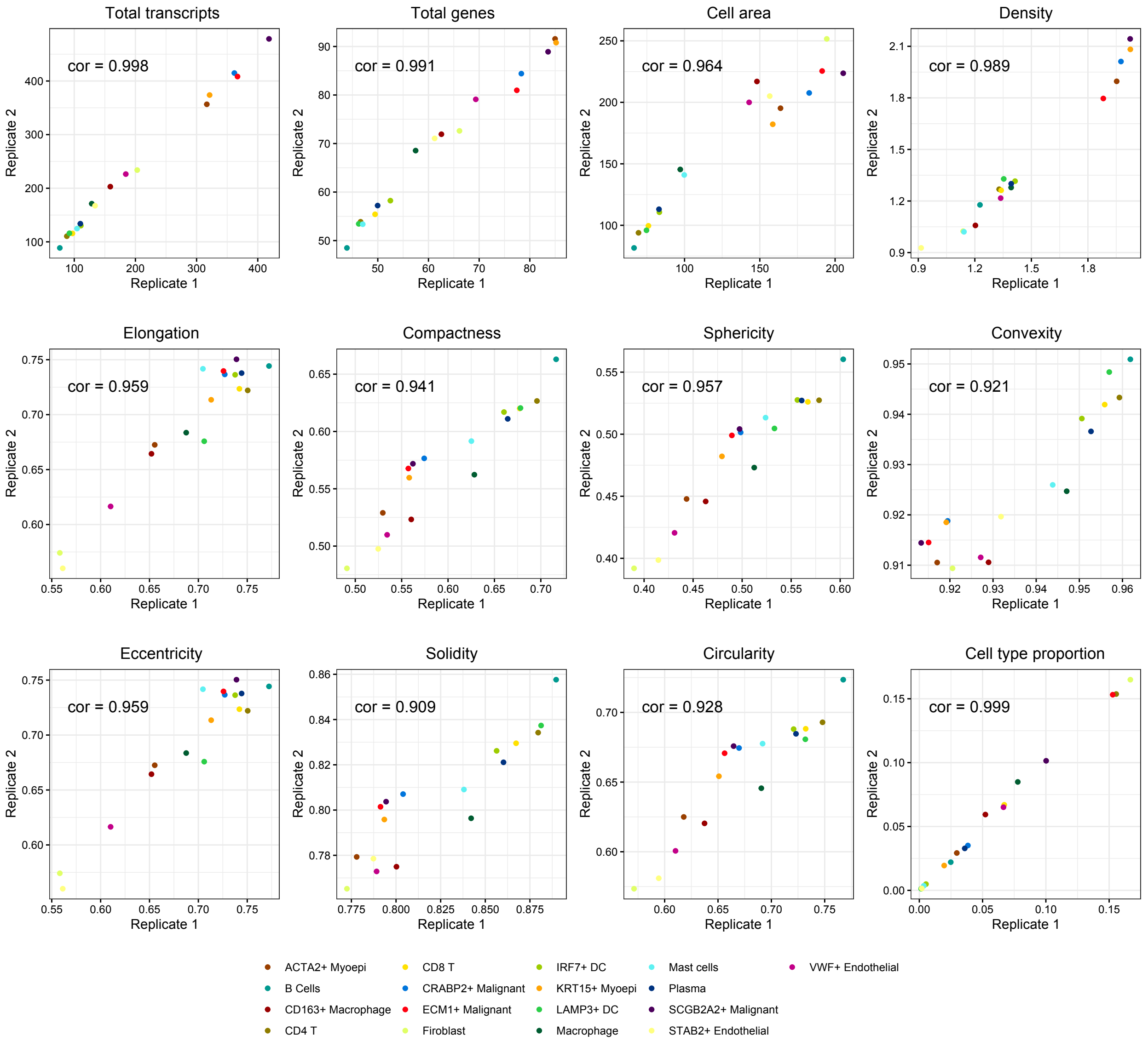


Supplementary Figure 11: Consistency between two replicates. A 3 × 4 scatter plots showing average cell-level baseline metrics, cell morphology metrics, and cell type proportions for cell types between two replicates.

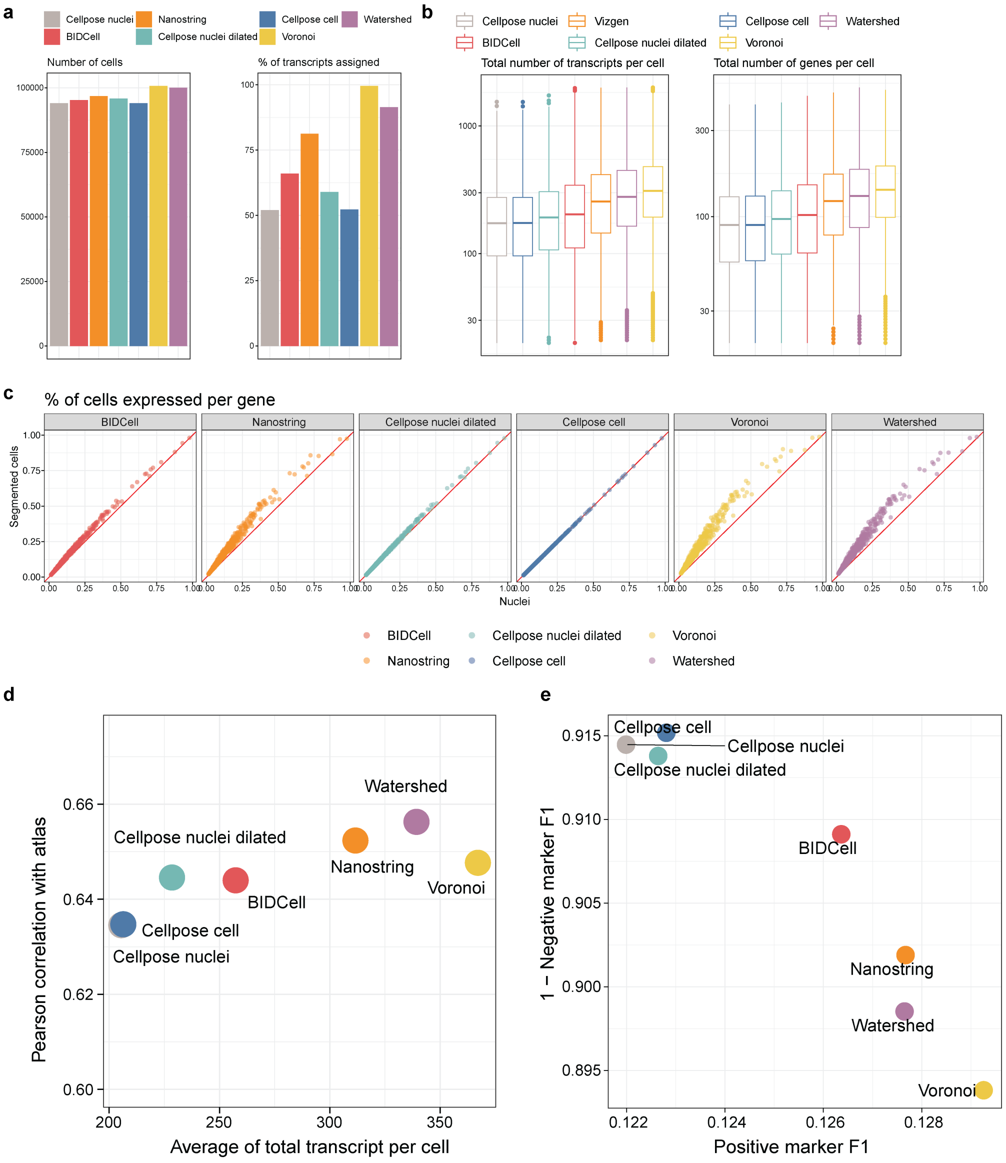


Supplementary Figure 12: Benchmarking results for CosMx-Lung. (a) Bar plot demonstrating overall characteristics, where the left panel shows the number of cells and the right panel shows the number of transcripts for each of the 7 methods. (b) Boxplot of cell-level quality metrics with total number of transcripts (left panel) and total number of genes (right panel). (c) Gene-level quality metric represented by a scatter plot of percentage of cells expressed for each gene between the nuclei vs. the cell body. (d) Scatter plot between correlation with Chromium expression (y-axis) and average total number of transcripts per cell (x-axis), where each dot represents a different method. (e) Scatter plot between positive marker F1 score and 1 - negative marker F1 for each method.

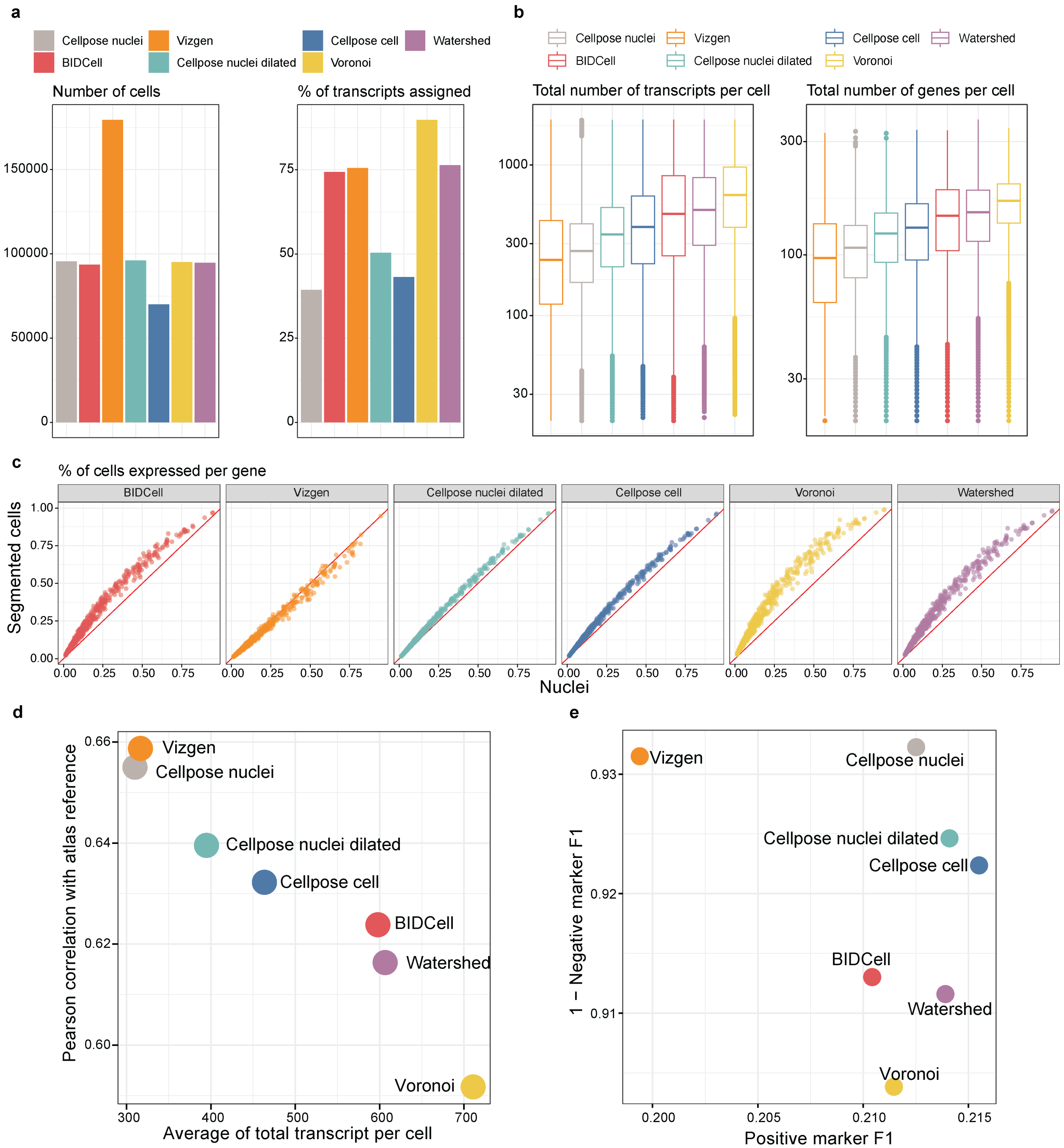


Supplementary Figure 13: Benchmarking results for MERSCOPE-Melanoma. (a) Bar plot demonstrating overall characteristics, where the left panel shows the number of cells and the right panel shows the number of transcripts for each of the 7 methods. (b) Boxplot of cell-level quality metrics with total number of transcripts (left panel) and total number of genes (right panel). (c) Gene-level quality metric represented by a scatter plot of percentage of cells expressed for each gene between the nuclei vs. the cell body. (d) Scatter plot between correlation with Chromium expression (y-axis) and average total number of transcripts per cell (x-axis), where each dot represents a different method. (e) Scatter plot between positive marker F1 score and 1 - negative marker F1 for each method.

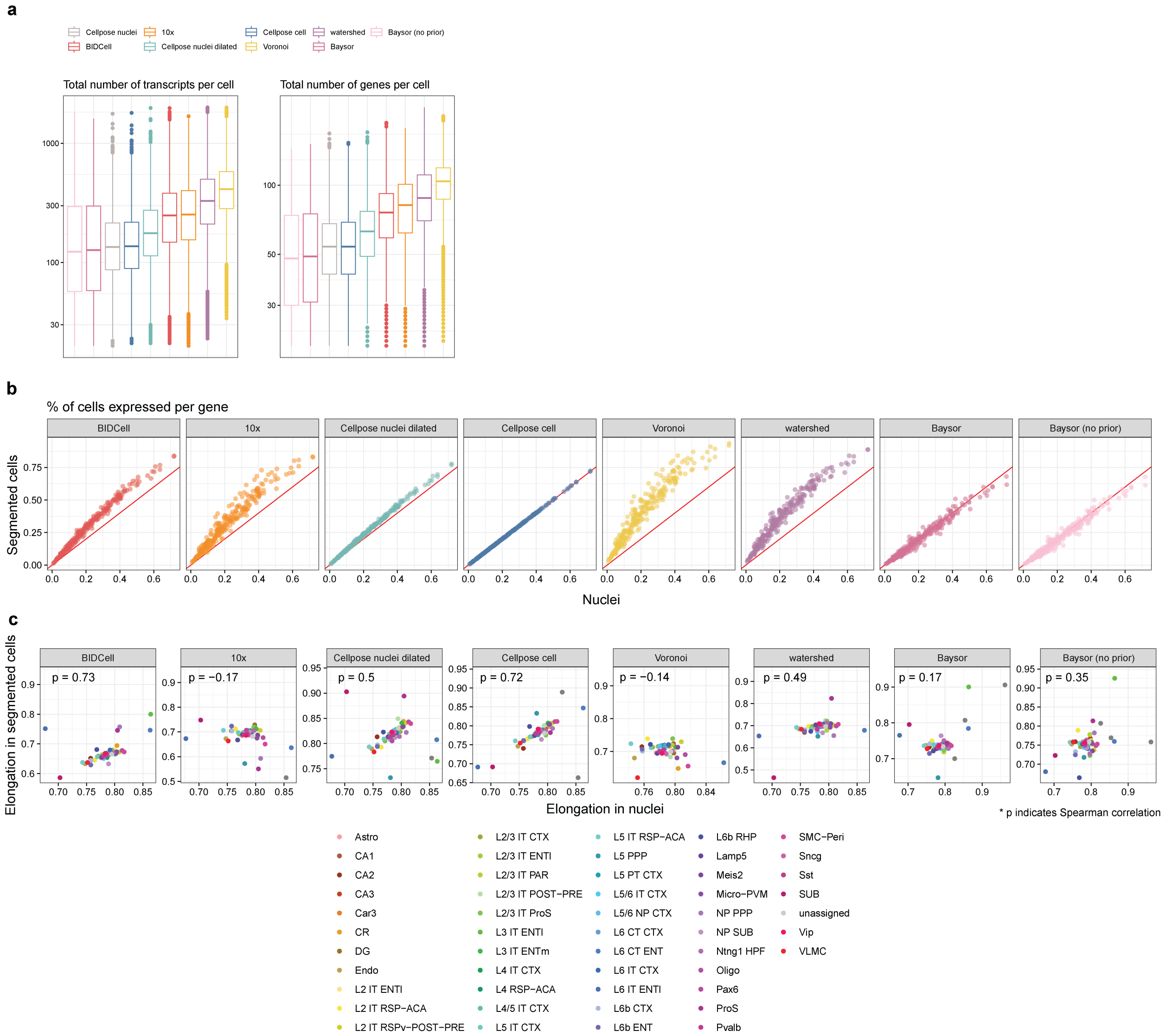


Supplementary Figure 14: Benchmarking results for Xenium-brain data. (a) Boxplot of cell-level quality metrics with total number of transcripts (left panel) and total number of genes (right panel). (b) Gene-level quality metric represented by a scatter plot of percentage of cells expressed for each gene between the nuclei vs. the cell body. (c) Cell morphology metrics represented by the elongation values between the nuclei and segmented cell, where each dot represents the average elongation metrics for each cell type.

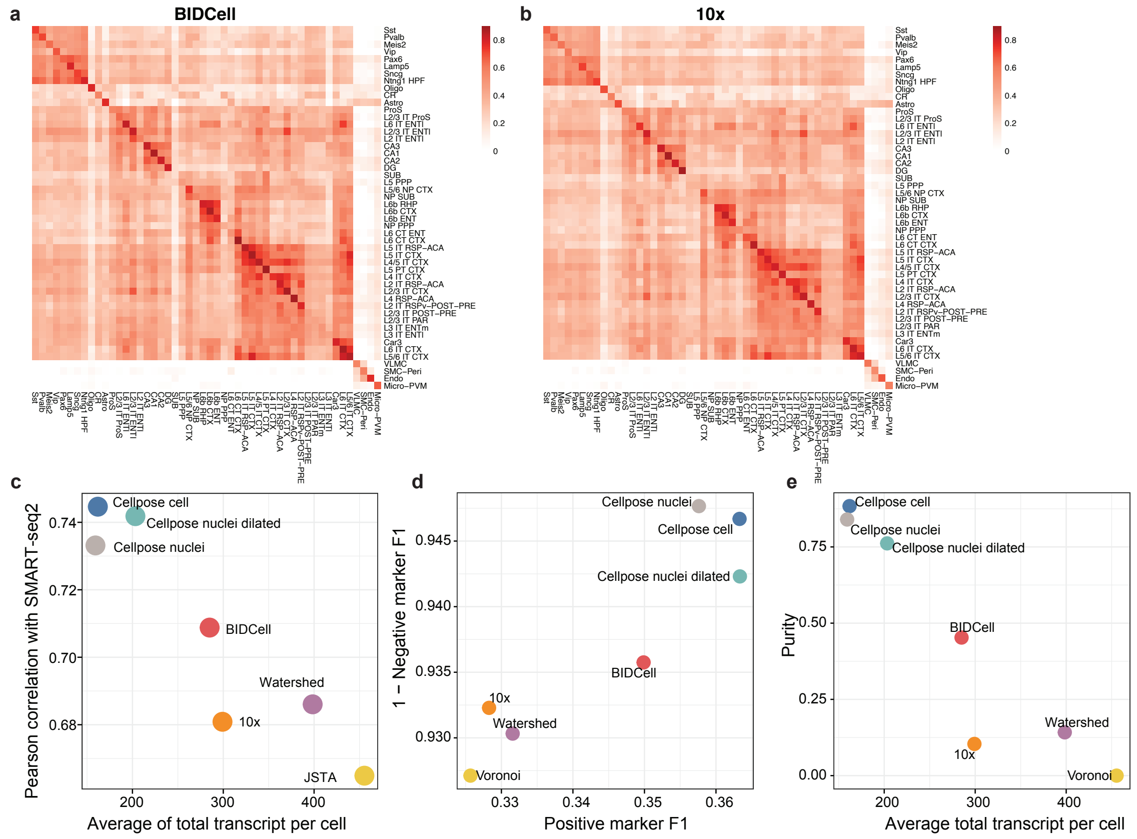


Supplementary Figure 15: Benchmarking results for Xenium-brain data. (a-b) Correlation heatmap of average expression between expression from Allan brain SMART-seq2 data (x-axis) and segmented cells from (a) BIDCell (y-axis); (b) 10x. (c) Scatter plot between correlation with Chromium expression (y-axis) and average total number of transcripts per cell (x-axis), where each dot represents a different method. (d) Scatter plot between positive marker F1 score and 1 - negative marker F1 for each method. (e) Scatter plot showing the purity F1 score against the average total transcripts per cell for each method.

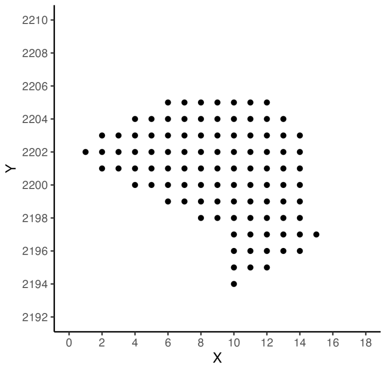


(a) Cell shape

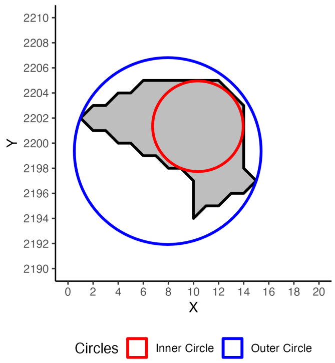


(b) Inscribing and circumscribing circles

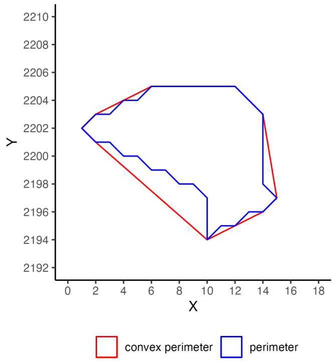


(c) Perimeter and convex perimeter

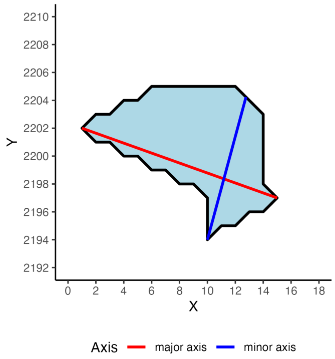


(d) Major and minor axis

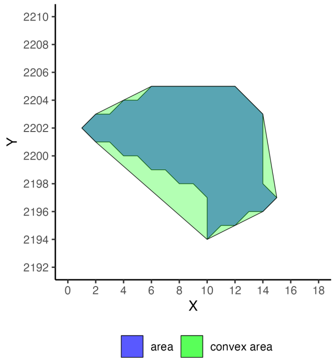


(e) Area and convex area

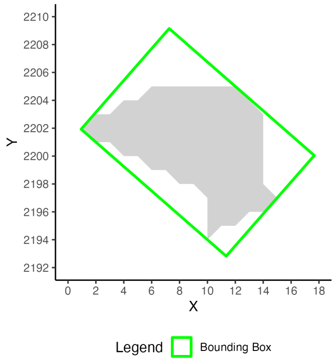


(f) Bounding box

Supplementary Figure 16: Cell metrics.
